# A highly polymorphic effector protein promotes fungal virulence through suppression of plant-associated Actinobacteria

**DOI:** 10.1101/2022.08.22.504754

**Authors:** Nick C. Snelders, Jordi C. Boshoven, Yin Song, Natalie Schmitz, Gabriel L. Fiorin, Hanna Rovenich, Grardy C.M. van den Berg, David E. Torres, Luigi Faino, Michael F. Seidl, Bart P.H.J. Thomma

**Affiliations:** University of Cologne, Institute for Plant Sciences, Cluster of Excellence on Plant Sciences (CEPLAS), 50674 Cologne, Germany; University of Utrecht, Theoretical Biology and Bioinformatics Group, Department of Biology, 3584CH Utrecht, The Netherlands; Wageningen University and Research, Laboratory of Phytopathology, Droevendaalsesteeg 1, 6708PB Wageningen, The Netherlands; State Key Laboratory of Crop Stress Biology for Arid Areas and College of Agronomy, Northwest A&F University, Yangling 712100, Shaanxi, China; Sapienza University of Rome, Department of Ambiental Biology, 00185 Rome, Italy

**Keywords:** Antimicrobial, avirulence factor, effector, holobiont, immune receptor, microbiota, pathogen

## Abstract

Plant pathogens secrete effector proteins to support host colonization through a wide range of molecular mechanisms, while plant immune systems evolved receptors to recognize effectors or their activities to mount immune responses to halt pathogens. Importantly, plants do not act as single organisms, but rather as holobionts that actively shape their microbiota as a determinant of health, and may thus be targeted by pathogen effectors as such. The soil-borne fungal pathogen *Verticillium dahliae* was recently demonstrated to exploit the VdAve1 effector to manipulate the host microbiota to promote vascular wilt disease in absence of the corresponding immune receptor Ve1. We now identified a multiallelic *V. dahliae* gene displaying ~65% sequence similarity to *VdAve1*, named *VdAve1-like* (*VdAve1L*). Interestingly, *VdAve1L* shows extreme sequence variation, including alleles that encode dysfunctional proteins, indicative of selection pressure to overcome host recognition. We show that the orphan cell surface receptor Ve2, encoded at the *Ve1* locus, does not recognize VdAve1L. Furthermore, we show that the full-length variant VdAve1L2 possesses antimicrobial activity, like VdAve1, yet with a divergent activity spectrum. Altogether, VdAve1L2 is exploited by *V. dahliae* to mediate tomato colonization through the direct suppression of antagonistic Actinobacteria in the host microbiota. Our findings open up strategies for more targeted biocontrol against microbial plant pathogens.

## INTRODUCTION

The plethora of microbes a plant associates with, its so-called microbiota, encompasses a diversity of microbes that establish a spectrum of symbiotic relationships with their host, ranging from commensalistic through endophytic to mutualistic and pathogenic (Hassani *et al*., 2019). To survey their microbiota for potentially pathogenic invaders, plants evolved complex immune systems comprising various types of receptors that betray microbial ingress, for instance through the recognition of microbe-associated molecular patterns (MAMPs) or microbially-secreted effector proteins (Chisholm *et al*., 2006; Jones & Dangl, 2006; Dodds & Rathjen, 2010; Thomma *et al*., 2011; Cook *et al*., 2015). If not recognized, microbial effectors play crucial roles during plant colonization. While they can benefit microbes in many ways, most of the effectors functionally characterized to date have been implicated in deregulation of host immune responses or other processes in host physiology (Rovenich *et al*., 2014; Cook *et al*., 2015; He *et al*., 2020; Wang *et al*., 2022). However, novel effector functions are still discovered. For instance, we recently demonstrated that some effectors are secreted to manipulate plant microbiota compositions to promote host colonization (Snelders *et al*., 2020, 2021, 2022).

Effector recognition in plants is mediated by resistance (*R*) genes, typically encoding receptors that reside on the cell surface or in the host cell cytoplasm, that detect effectors or their activities to activate effector-triggered immunity (ETI), often leading to avirulence of the pathogen (Chisholm *et al*., 2006; Jones & Dangl, 2006; Dodds & Rathjen, 2010; Thomma *et al*., 2011; Cook *et al*., 2015). Consequently, effectors that are recognized by R proteins are referred to as <u>av</u>i<u>r</u>ulence factors (Avr)(Li *et al*., 2020). To evade ETI and restore the ability to colonize their hosts, pathogens are known to inactivate, purge or mutate their avirulence genes, or to evolve novel effectors that suppress ETI (van Kan *et al*., 1991; Armstrong *et al*., 2005; Gout *et al*., 2007; Stergiopoulos *et al*., 2007; Zhou *et al*., 2013; Wu *et al*., 2014; Niu *et al*.,2016; Schmidt *et al*., 2016; Praz *et al*., 2017).

*Verticillium dahliae* is a soil-borne fungal pathogen that causes vascular wilt disease in hundreds of plant species (Fradin & Thomma, 2006). The presumed asexual fungus generates genomic diversity through extensive chromosomal rearrangements and segmental duplications that gave rise to dynamic so-called lineage-specific (LS) regions, more recently referred to as <u>a</u>daptive <u>g</u>enomic <u>r</u>egions (AGRs) (Klosterman *et al*., 2011; de Jonge *et al*., 2013; Faino *et al*.,2016; Shi-Kunne *et al*., 2018; Cook *et al*., 2020). These AGRs display extensive presence/absence variation (PAV) between *V. dahliae* strains, are rich in repeats and transposable elements, and have a distinct chromatin profile (Klosterman *et al*., 2011; de Jonge *et al*., 2013; Faino *et al*., 2016; Cook *et al*., 2020; Torres *et al*., 2021). Additionally, AGRs are enriched in *in planta-induced* genes and harbor effector genes that are crucial for disease establishment (de Jonge *et al*., 2012, 2013; Kombrink *et al*., 2017). Thus, like other filamentous plant pathogens, *V. dahliae* possesses a compartmentalized genome in which (a)virulence factors locate in regions of increased plasticity when compared with the core genome, an arrangement often referred to as a “two-speed” genome (Raffaele & Kamoun, 2012; Dong *et al*., 2015; Torres *et al*., 2020). Intriguingly, *V. dahliae* AGRs that are conserved between strains display enhanced sequence conservation when compared with core genomic regions (Depotter *et al*., 2019), underscoring that accelerated evolution in these regions is predominantly mediated by presence–absence polymorphisms.

Only few genetic resistance sources to *V. dahliae* have been identified. In tomato, the *Ve* locus provides resistance against *V. dahliae* and has been introgressed into most commercial tomato cultivars (Schaible *et al*., 1951; Diwan *et al*., 1999). The *Ve* locus contains two closely linked genes, *SlVe1* and *SlVe2*, that both encode extracellular leucine-rich repeat receptor-like proteins (eLRR-RLPs), of which only SlVe1 was confirmed to confer *V. dahliae* resistance (Diwan *et al*., 1999; Kawchuk *et al*., 2001; Fradin *et al*., 2009). Since its deployment in the 1950s, resistance-breaking strains appeared that have been assigned to race 2, whereas strains that are contained belong to race 1 (Alexander, 1962). Recently, the single dominant *SlV2* locus was shown to mediate race 2 resistance in *Solanum neorickii* and was introgressed in particular tomato rootstock cultivars (Usami *et al*., 2017). However, resistance-breaking strains already appeared, forming race 3 (Usami *et al*., 2017; Chavarro-Carrero *et al*., 2021). Comparative *V. dahliae* population genomics led to the identification of the avirulence factors corresponding to *SlVe* and *SlV2* resistance, namely *VdAve1* and *VdAv2*, respectively (de Jonge *et al*., 2012; Chavarro-Carrero *et al*., 2021). Both effector genes are located in AGRs and display PAV between *V. dahliae* strains. While the molecular function of *VdAv2* remains unknown, we demonstrated that VdAve1 promotes virulence of *V. dahliae* on host plants lacking SlVe1 through selective antimicrobial activity to manipulate host microbiota compositions. More specifically, we showed that VdAve1 facilitates *V. dahliae* colonization of tomato and cotton through the direct suppression of associated antagonistic bacteria of the Sphingomonadales order (Snelders *et al*., 2020). Intriguingly, race 1 strains of *V. dahliae* do not display any allelic variation for *VdAve1*, and only two allelic *VdAv2* variants have been identified among race 2 strains that differ by one non-synonymous single nucleotide polymorphism (SNP) and that both activate *Av2*-mediated immunity (Chavarro-Carrero *et al*., 2021). Hence, to date, *V. dahliae* has exclusively been described to evade ETI through loss of complete avirulence genes, and not through gene inactivation or the evolution of allelic effector variants.

## MATERIALS AND METHODS

### *Verticillium dahliae* genomics

Genome sequence data used in this study was obtained previously (Klosterman *et al*.,2011; de Jonge *et al*., 2012, 2013; Faino *et al*., 2015; Chavarro-Carrero *et al*., 2021). BLAST searches were performed with BLAST+ version 2.11.0 using standard parameters and the nucleotide sequences of *VdAve1* or each of the *VdAve1L* alleles as query. Strains lacking *VdAve1L* in their genome assemblies were further assessed for the presence of the gene by PCR using the primers listed in Supplementary Table 1. Amplicons obtained from each strain were sequenced for allele determination. Discontinuity in the *VdAve1L3* allele was identified by mapping the genomic paired-end sequences to the *V. dahliae* strain JR2 reference genome using BWA-mem (Li & Durbin, 2009; Faino *et al*., 2015). The genetic diversity and the population structure of the sequenced *V. dahliae* strains was assessed using the reference sequence alignment-based phylogeny builder REALPHY (version 1.12) (Bertels *et al*., 2014) and Bowtie2 (version 2.2.6) (Langmead & Salzberg, 2012) to map genomic reads against the *V. dahliae* strain JR2 gapless genome. A maximum likelihood phylogenetic tree was built using RAxML (version 8.2.8) (Stamatakis, 2014).

### Presence-absence variation (PAV) analysis

PAV was identified using whole-genome alignments of 52 *V. dahliae* strains. Paired-end short sequencing reads were mapped to reference *V. dahliae* strain JR2 (Faino *et al*., 2015) using BWA-mem with default settings (Li & Durbin, 2009). Long-reads were mapped to *V. dahliae* JR2 using minimap2 with default settings (Li, 2018). Using the Picard toolkit (http://broadinstitute.github.io/picard/), library artifacts were marked and removed with - *MarkDuplicates* followed by *-SortSam* to sort the reads. Raw read coverage was averaged per 100 bp non-overlapping windows using the *-multicov* function of BEDtools (Quinlan & Hall, 2010). Then, raw read coverage values were transformed to a binary matrix by applying a cuto-ff of 10 reads for short-read data; >=10 reads indicate presence (1) and <10 reads indicate absence (0) of the respective genomic region. In the case of long-read data, a cut-off of 1 read was applied; >=1 reads indicate presence (1) and <1 reads indicate absence (0). This matrix was further summarized to obtain the total number of presence/absence counts in each 100 bp genomic window within 50 kb upstream and downstream of the *VdAve1L* locus.

### SNP rate determination of *V. dahliae* genes

To assess if the *VdAve1L* allele indeed displays an unusual degree of sequence variation, we determined the number of SNPs in the CDS of all genes in the genome of *V. dahliae* strain JR2 compared to 41 other strains that we previously sequenced using the Illumina platform (Torres *et al*., 2021). Next, we normalized the identified number of SNPs based on the length of the corresponding CDS to determine a SNP rate.

### *VdAve1L2* gene expression

To determine *in planta* expression of *VdAve1L2*, tomato plants (*Solanum lycopersicum*)cultivar MoneyMaker were inoculated with *V. dahliae* strain DVD-S26 as previously described (Fradin *et al*., 2009). Stems of five mock and five inoculated plants were harvested at 7, 14 and 21 days after inoculation. *In vitro* expression of *VdAve1L2* was assessed in mycelium of *V. dahliae* strain DVD-S26 grown for 5 days in triplicate on PDA plates and in soil extract (Snelders *et al*., 2020). Total RNA of all samples was extracted using the Maxwell76 LEV Plant RNA kit (Promega, Leiden, the Netherlands) and cDNA was synthesized using the M-MLV Reverse Transcriptase (Promega, Leiden, the Netherlands). Real-time PCR was conducted using a C1000 Touch™ Thermal Cycler (Bio-Rad, California, USA) and the qPCR SensiMix kit (BioLine, GC Biotech BV, Alphen aan den Rijn, The Netherlands) using the primers listed in Supplementary Table 1. Real-time PCR conditions were as follows: an initial 95°C denaturation step for 10 minutes followed by denaturation for 15 seconds at 95°C, annealing for 60 seconds at 60°C, and extension at 72°C for 40 cycles.

### Generation of *V. dahliae* mutants

To generate *VdAve1L2* deletion lines, primers were designed to amplify approximately 1500 bp up- and downstream of the *VdAve1L2* CDS (Supplementary Table 1). Both amplicons were used to generate a USER-friendly cloning construct to replace *VdAve1L2* by a hygromycin cassette through homologous recombination (Frandsen *et al*., 2008). To complement the *VdAve1L2* deletion mutant, a PCR fragment was amplified from genomic DNA containing the complete *VdAve1L2* CDS and approximately 1000 bp up- and downstream sequences (Supplementary Table 1) and cloned into the binary vector pCG (Zhou *et al*., 2013). To generate the *V. dahliae* DVDS-S26 mutants expressing *VdAve1L2* under control of the *VdAve1* promoter, the coding sequence of *VdAve1L2* was amplified and cloned into pFBT005. *V. dahliae* transformations were performed as described previously (Santhanam, 2012).

### Disease assays

Inoculation of tomato plants to determine the virulence of the *V. dahliae* was performed as described previously (Fradin *et al*., 2009). Accumulation of *V. dahliae* biomass in the tomato plants was quantified with real-time PCR on the genomic DNA by targeting the internal transcribed spacer (ITS) region of the ribosomal DNA using the primers listed in Supplementary Table 1. To assess the importance of the suppression of Actinobacteria by VdAve1L2 for tomato colonization by *V. dahliae*, we germinated tomato MoneyMaker seeds in a sealed beaker on sterile potting soil (Lentse potgrond) supplemented with water-treated tomato root microbiota or vancomycin-treated tomato root microbiota. Vancomycin-treatment of the tomato root microbiota was performed as described previously, with slight modifications (Lee *et al*., 2021). Briefly, roots with rhizosphere soil were harvested from six-week-old tomato MoneyMaker plants grown on potting soil (Lentse potgrond). The material collected from four plants was pooled and ground to a fine powder in liquid nitrogen using mortar and pestle. Subsequently, the ground material was split and transferred to 300 mL 2.5 mM MES pH 6.0 supplemented with 500 μg/mL vancomycin or water. The suspensions were incubated for 3 hours at 30°C and 120 rpm. Finally, the suspensions were divided in fractions of 50 mL, snap frozen and stored at −20°C until use. Tomato seeds were surface-sterilized by incubation for 5 min in 2% sodium hypochlorite. Next, the surface-sterilized tomato seeds were washed three times using sterile water and transferred to sealed beakers containing 60 grams of sterilized potting soil and 50 mL of the water-treated or vancomycin-treated microbial suspensions. After 14 days, seedlings colonized by the microbial communities were root dipped in spore suspensions of *V. dahliae* strain DVD-S26, a *VdAve1L2* deletion mutant or water (mock) as described previously (Fradin *et al*., 2009) and transferred to pots containing sterile river sand soaked in Hoagland nutrient solution.

### *Agrobacterium tumefaciens-mediated* transient expression in *N. tabacum*

*Nicotiana tabacum* cv. Petite Havana SR1 was infiltrated with GV3101 *A. tumefaciens* strains carrying pSOL2092 constructs for expression of *Ve1* or *Ve2* and an *VdAve1* (-like) construct, as described previously (Zhang *et al*., 2013). Plants were transferred to a climate chamber and incubated at 22°C and 19°C during 16-h day and 8-h night periods, respectively, with 70% relative humidity. Leaves were inspected for HR at 5 dpi.

### Protein production, purification and refolding

Heterologous production and purification of VdAve1L2 was performed as described previously for VdAve1 (Snelders *et al*., 2020).

### Oxford Nanopore Technology sequencing

Library preparation of the PCR fragment was performed according to the protocol of Oxford Nanopore, skipping the DNA fragmentation step. The library was loaded on a Nanopore flow cell. The run yielded about 870 high quality long reads with an average length of 3,869 bp with the longest read being ~14 kb. Using Nanocorrect, we used all the obtained reads to correct the 50 longest reads, of which 28 were corrected to generate a consensus. Finally, reads were used for BLAST analysis to confirm the presence of *VdAve1L3* fragments at both ends.

### Root microbiota analysis

Tomato inoculations were performed as described previously (Fradin *et al*., 2009). After 10 days, plants were carefully uprooted and gently shaken to remove loosely adhering soil from the roots. Next, roots with rhizosphere soil from two tomato plants were pooled to form a single biological replicate. Alternatively, the tomato plants that received the water-treated and vancomycin-treated microbial communities where uprooted 18 days post inoculation with *V. dahliae* and a single root system with adhering river sand was collected as a biological control. All samples were flash-frozen in liquid nitrogen and ground using mortar and pestle. Genomic DNA isolation was performed using the DNeasy PowerSoil Kit (Qiagen, Venlo, The Netherlands. Sequence libraries were prepared following amplification of the V4 region of the bacterial 16S rDNA (341F and 785R) and paired ends (300 bp) were sequenced using the MiSeq sequencing platform (Illumina) at Baseclear (Leiden, The Netherlands). Data analyses were performed as described previously (Snelders *et al*., 2020).

### Bacterial isolates

Bacterial strains *B. subtilis* AC95, *S. xylosus*. M3, *P. corrugata* C26 and *Ralstonia* sp. M21 were obtained from our in house endophyte culture collection (Snelders *et al*., 2020). Bacterial strains *Aeromicrobium sp*. (DSM 102283), *C. chitinilytica* (DSM 17922), *F. peucedani* (DSM 22180), *J. huperziae* (DSM 46866), *Leifsonia sp*. (DSM 102435) and *N. plantarum* (DSM 11054) were obtained from the DSMZ culture collection (Braunschweig, Germany). Bacterial strains *Novosphingobium* sp. A (NCCB 100261), *S. macrogoltabida* (NCCB 95163) and *Sphingobacterium* sp. (NCCB 100093) were obtained from the Westerdijk Fungal Biodiversity Institute (Utrecht, the Netherlands).

### Antimicrobial activity assays

*In vitro* antimicrobial activity assays were performed as described previously using 0.2x tryptone soy broth as growth medium for the bacteria (Snelders *et al*., 2020).

### *In vitro* competition assays

Following eleven days of cultivation on PDA, the conidiospores of *V. dahliae* strains DVDS26 *ΔVdAve1L2*, DVD-S26 *ΔVdAve1L2* + *pVdAve1::VdAve1L2* #1 and DVD-S26 *ΔVdAve1L2* + *pVdAve1::VdAve1L2* #2 were harvested from plate and stored at −80° C at a concentration of 4*10^5^ spores/mL in low salt TSB (17 g/L tryptone, 3 g/L soy peptone, 0.5 g/L NaCl, 2.5 g/L K2HPO4 and 2.5 g/L glucose) supplemented with 10% glycerol until use. Next, bacterial isolates were grown on low salt TSA at 28°C. Single colonies were selected and grown overnight at 28°C while shaking at 150 rpm. Overnight cultures were resuspended to an OD_600_=0.02 in fresh low salt TSB, while the fungal spore suspensions were allowed to thaw at room temperature. Finally, the bacterial and fungal spore suspensions were mixed in 500 μl of low salt TSB to a final concentration of OD_600_=0.01 and 10^3^ spores/mL, respectively.

Following six days of incubation at 22°C, the microbial suspensions were transferred to clear 24-well flat-bottom polystyrene tissue culture plates to allow imaging of fungal growth using an SZX10 stereo microscope (Olympus) equipped with a EP50 camera (Olympus).

## RESULTS

### Identification of the highly polymorphic *VdAve1-like* gene in an adaptive genomic region of *Verticillium dahliae*

Using the gapless genome assembly of *V. dahliae* strain JR2 (Faino *et al*., 2015), a similarity search with the coding sequence (CDS) of *VdAve1* revealed a gene with 67% nucleotide similarity that we further refer to as *VdAve1L* (for *VdAve1-like*). However, while *VdAve1* encodes a 134 amino acid protein (de Jonge *et al*., 2012), *VdAve1L* only encodes a 24 amino acid protein due to a premature stop codon (Fig. 1a). Searches in the genomes of 51 additional *V. dahliae* isolates (Klosterman *et al*., 2011; de Jonge *et al*., 2012; Faino *et al*., 2015; Chavarro-Carrero *et al*., 2021) identified the gene in 42 isolates (Fig. 1a,b; Supplementary Fig. 1), albeit with considerable allelic variation. In total we identified six allelic variants (*VdAve1L1* to *VdAve1L6*) that share 93-99% sequence similarity (Fig. 1a,b). Like *VdAve1L1*, also *VdAve1L3* and *VdAve1L4* encode truncated 24 amino acid proteins (Fig. 1a). In contrast, and similar to *VdAve1, VdAve1L2* and *VdAve1L5* encode 134 amino acid proteins including an 18 amino acid N-terminal signal peptide (Fig. 1a). Finally, VdAve1L6 only differs by one amino acid when compared with VdAve1L5 and is truncated after 120 amino acids (Fig. 1a).

**Figure 1.**
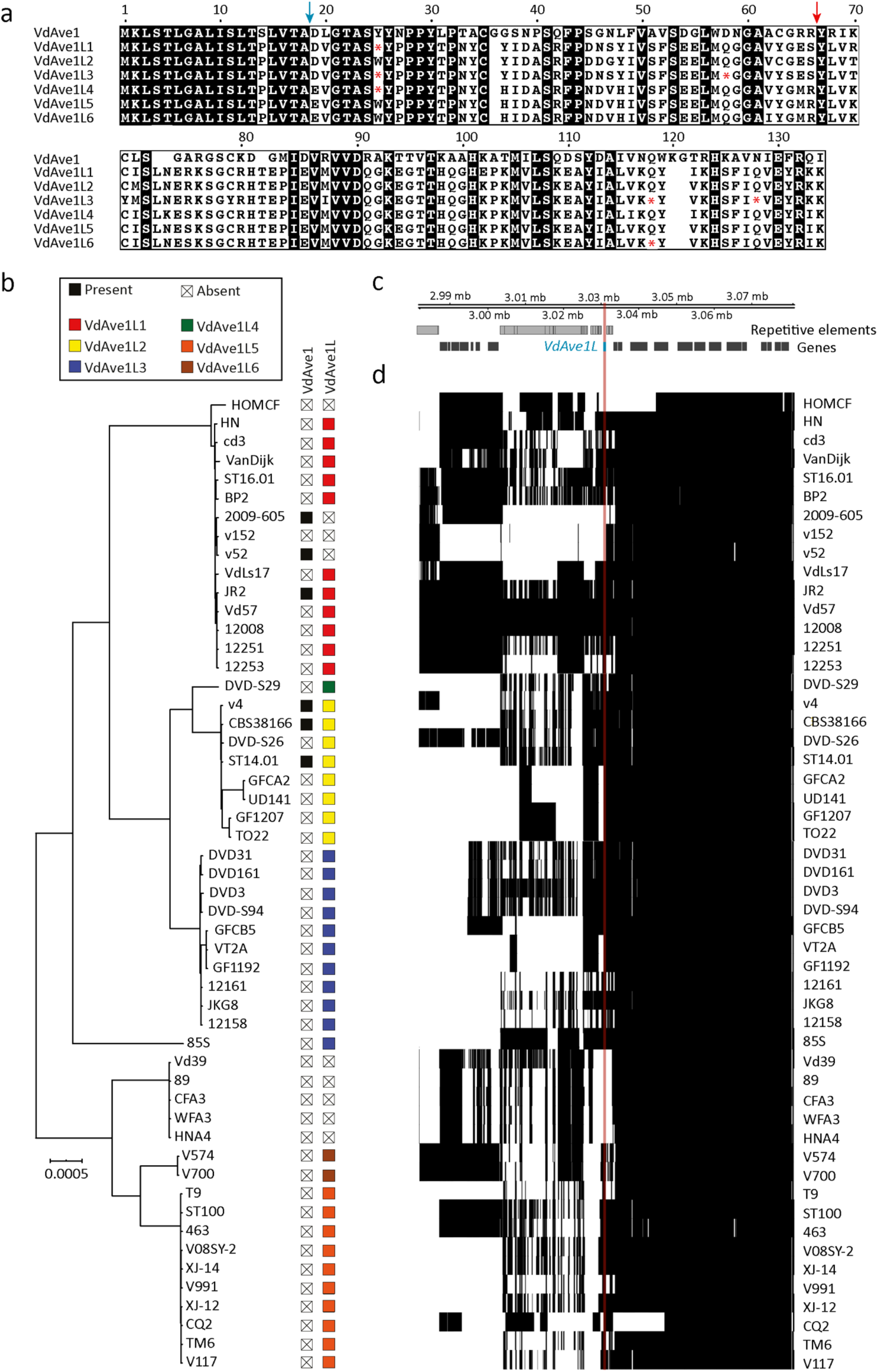
*Verticillium dahliae* strains carry a multiallelic *VdAve1-like* gene. **(a)** Protein sequence alignment of the identified VdAve1L proteins and VdAve1. Red asterisks (*) indicate stop codons in the *VdAve1L* alleles. The blue arrow indicates the predicted signal peptide cleavage site, and the red arrow indicates the site where discontinuity is observed in *VdAve1L3*. **(b)** Phylogenetic tree of sequenced *V. dahliae* strains and the presence-absence variation of *VdAve1* and *VdAve1L* alleles in the corresponding genomes. **(c-d)** *VdAve1L* is located in an adaptive genomic region of the *V. dahliae* genome. The matrix displays the presence-absence (black/white) variation in 100 bp non-overlapping windows in the genomic region surrouding *VdAve1L*. The upper panel displays the annotated genes and repetitive elements identified in this genomic region, indicated in black and grey, respectively.

Genome assemblies consistently assigned sections of *VdAve1L3* on separate contigs, suggesting discontinuity of this allele. (Supplementary Fig. 2a), which was further supported by PCR analysis on strains DVD-3, DVD-31, DVD-S94 and DVD-161 (Supplementary Fig. 2b). To investigate whether the discontinuity is caused by a chromosomal rearrangement or transposable element insertion, we performed the PCR on strain DVD-3 with prolonged elongation time, yielding an amplicon of ~7 kb (Supplementary Fig. 2c). Subsequent sequence analysis revealed that *VdAve1L3* is interrupted by a long terminal repeat retrotransposon classified as VdLTRE3 (Faino *et al*., 2016).

As expected based on the observed PAV among some *V. dahliae* strains (Fig. 1b), *VdAve1L* is localized in an AGR (Fig 1c,d) (Cook *et al*., 2020). However, the allelic variation of *VdAve1L* is unexpected given the previously observed commonly increased sequence conservation of AGR sequences that are shared among *V. dahliae* strains (Depotter *et al*., 2019). Intriguingly, further analysis revealed that *VdAve1L* displays the highest SNP rate of all genes in the *V. dahliae* genome (Supplementary Fig. 3), and that all of the identified SNPs, 30 in total (Supplementary Fig. 4), cause protein sequence variation. Thus, *VdAve1L* is a highly polymorphic gene that displays accelerated evolution by PAV, transposon-mediated sequence disruption, and sequence variation.

### VdAve1L proteins are not recognized by SlVe2

The extreme sequence variation of VdAve1L is likely the result of selection pressure to overcome recognition by a plant immune receptor. The only *R* gene known to confer resistance to *V. dahliae* is tomato *SlVe1* (Kawchuk *et al*., 2001; Fradin *et al*., 2009), which resides in the *Ve* locus together with *SlVe2* that similarly encodes a receptor-like protein (Fradin *et al*., 2009). *SlVe2* is expressed similarly as *SlVe1*, yet the encoded receptor does not recognize VdAve1 and its function remains unknown (Fradin *et al*., 2009; de Jonge *et al*., 2012). Thus, we tested if SlVe2 can recognize a current or predicted ancestral VdAve1L variant. To this end, we reversed disruptive mutations in *VdAve1L1, VdAve1L3* and *VdAve1L4* by replacing premature stop codons with corresponding codons in *VdAve1L2* and *VdAve1L5* to yield the derivatives *VdAve1L1*, VdAve1L3\** and *VdAve1L4\** (Fig. 2a,b). Similarly, we replaced stop codons in *VdAve1L3\** and removed the retrotransposon (Fig. 2a,b). Additionally, based on an alignment consensus sequence we constructed the predicted common *VdAve1L* ancestor *VdAve1L*\*\* (Fig.2a,b). Subsequently, we co-expressed the various genes with *SlVe2* in *Nicotiana tabacum*,and with *SlVe1* as a negative control, but no hypersensitive response (HR) could be observed except upon co-expression of *SlAve1* (Fig. 2c). Consequently, it is unlikely that SlVe2 recognized VdAve1L and drove its diversification.

**Figure 2.**
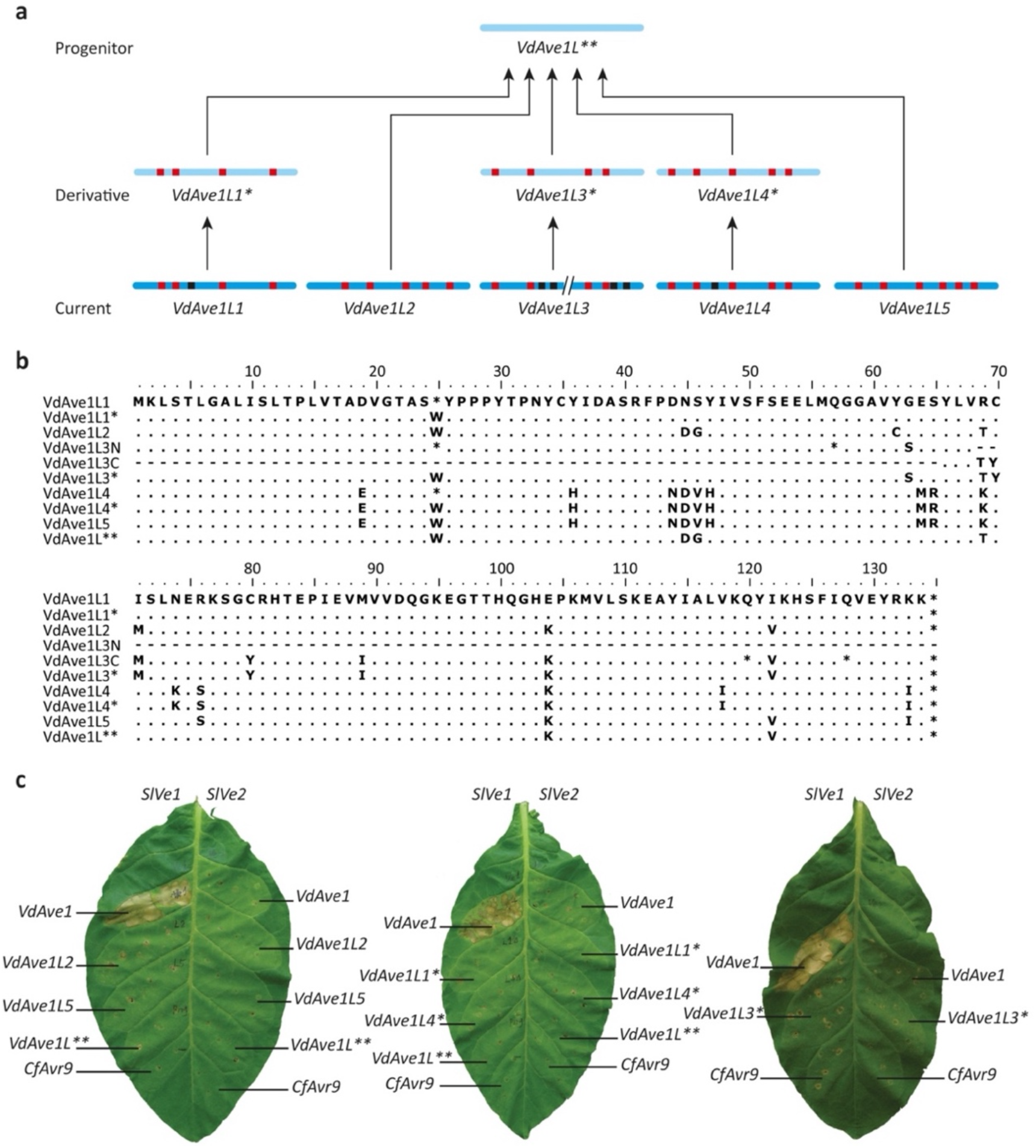
VdAve1L proteins are not recognized by the immune receptor SlVe2. **(a)** Schematic overview of the relationship between the currently identified *VdAve1L* alleles (excluding *VdAve1L6)*, the derived alleles *VdAve1L1*, VdAve1L3*, VdAve1L4\** and the putative progenitor allele (*VdAve1L*\*\*). The blue bars indicate the alleles, the line with arrows indicates the relationship between the current alleles, the artificial derivatives, and the progenitor. Within the blue bars the red blocks indicate point mutations, the black blocks indicate premature stop codons and the double slash (//) indicates discontinuity. **(b)** Alignment of VdAve1L proteins (VdAve1L1 – VdAve1L5), derivatives (VdAve1L1*, VdAve1L3* and VdAve1L4*), the progenitor (VdAve1L**), the N-terminal sequence of VdAve1L3 (VdAve1L3N) and the C-terminal sequence of VdAve1L3 (VdAve1L3C). Dots (.) indicate amino acid conservation, lines (-) indicate the absence of an amino acid, and an asterisk (*) indicates a stop codon at the respective position. **(c)** Co-expression of *VdAve1L* alleles with the tomato immune receptors *SlVe1* and *SlVe2*. Co-expression of *VdAve1* with *SlVe1* serves as positive control for the induction of a hypersensitive response. The sequence unrelated *Cladosporium fulvum* effector Avr9 serves as a negative control for recognition by SlVe1 and SlVe2.

### VdAve1L2 is a virulence factor that functionally diverged from VdAve1

Only the alleles *VdAve1L2* and *VdAve1L5* encode full length proteins (Fig. 1), yet none of the strains that encodes VdAve1L5 is pathogenic on tomato (Li, 2019). To determine if VdAve1L2, like VdAve1, contributes to virulence on tomato, we assessed its expression in *V. dahliae* race 2 strain DVD-S26 during host colonization, showing clear expression *in planta* (Supplementary Fig. 5). Interestingly, while we previously also detected expression of *VdAve1* during cultivation *in vitro* on PDA and in soil extract (Snelders *et al*., 2020), we did not detect *VdAve1L2* expression under these conditions (Supplementary Fig. 5).

To determine the importance of VdAve1L2 for tomato colonization, we generated deletion mutants in the race 2 strain DVD-S26 and in the race 1 strains ST14.01 and CBS38166 (Supplementary Fig. 6). Inoculation of tomato plants with the deletion mutants of the race 1 strains revealed no virulence contribution of VdAve1L2 (Fig 3a,b). Strikingly, however, we detected strongly compromised tomato colonization of the *VdAve1L2* deletion mutant in strain DVD-S26 (Fig 3c), which was restored in a complementation mutant (Fig. 3c). Thus, VdAve1L2 contributes to *V. dahliae* virulence on tomato in the absence of *VdAve1*. To address the hypothesis that VdAve1 and VdAve1L2 are functionally redundant, we introduced *VdAve1L2* under control of the *VdAve1* promoter in a *VdAve1* deletion mutant of the JR2 strain and tested its virulence on tomato, yet *VdAve1L2* failed to restore the virulence penalty of *VdAve1* deletion. Collectively, our data suggest that VdAve1 and VdAve1L2 have functionally diverged.

**Figure 3.**
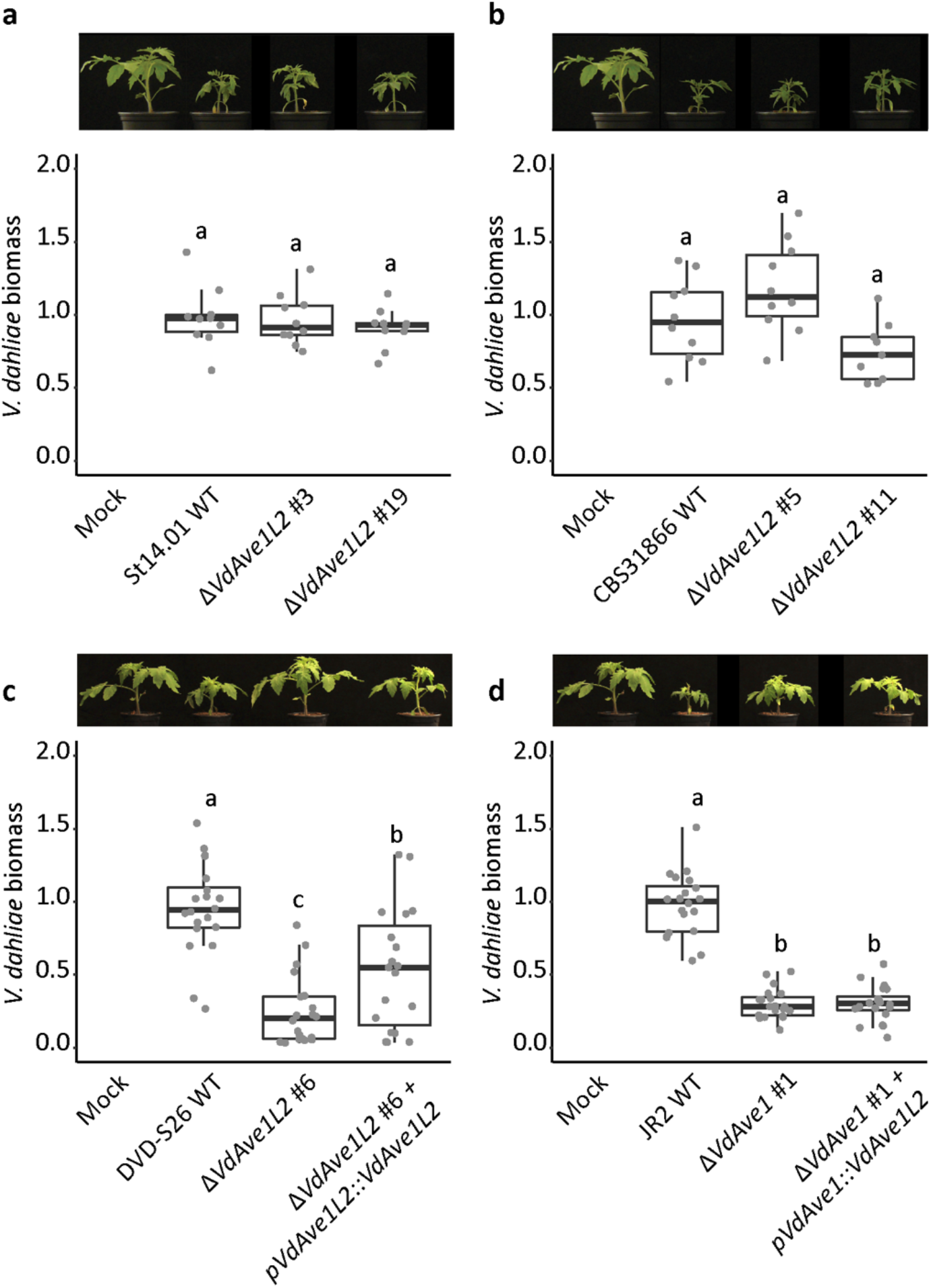
VdAve1L2 is a virulence factor of *V. dahliae* that functionally diverged from VdAve1. **(a-b)** VdAve1L2 does not contribute to virulence of *V. dahliae* race 1 strains St14.01 **(a)** and CBS31866 **(b)** on tomato. Photos display representative stunting symptoms of tomato plants 14 days post inoculation with wild-type *V. dahliae* strains and the corresponding *VdAve1L2* deletion mutants. *V. dahliae* biomass in tomato stems was quantified by real-time PCR. Letters represent non-significant biomass differences (one-way ANOVA and Tukey’s post hoc test; p<0.05; N=10). **(c-d)** VdAve1L2 contributes to virulence of *V. dahliae* race 2 strain DVD-S26 on tomato **(c)** but fails to restore the virulence that is lost by *V. dahliae* race 1 strain JR2 upon deletion of *VdAve1*. **(d)** Photos display representative stunting phenotypes of tomato plants 14 days post inoculation with the wild-type *V. dahliae* strains, the corresponding *VdAve1L2* or *VdAve1* deletion mutants, and the mutants expressing *VdAve1L2* under control of its native or *VdAve1* promoter. *V. dahliae* biomass in tomato stems was quantified by real-time PCR. Letters represent non-significant biomass differences (one-way ANOVA and Tukey’s post hoc test; p<0.05; N≥17).

### VdAve1L2 promotes *V. dahliae* virulence through suppression of Actinobacteria

While most effector proteins functionally characterized to date act in manipulation of host physiology, we recently showed that VdAve1 is an antibacterial effector protein that is secreted by *V. dahliae* to suppress microbial antagonists in the microbiomes of its hosts (Snelders *et al*., 2020). Thus, we hypothesized that VdAve1L2 may similarly exert antibacterial activity. *In vitro* assays previously revealed a strong activity of VdAve1 on the Gram positive bacterium *Bacillus subtilis* (Snelders *et al*., 2020). Interestingly, VdAve1L2 affected *B. subtilis* growth as well, albeit markedly less effectively (Fig. 4a). Furthermore, similar to VdAve1 (Snelders *et al*., 2020), VdAve1L2 inhibited the growth of plant-associated *Novosphingobium* sp. and *Staphylococcus xylosus*, but not of *Agrobacterium tumefaciens*, *Pseudomonas corrugata* and *Ralstonia* sp.. However, in contrast to VdAve1, VdAve1L2 did not inhibit growth of *Sphingobacterium* sp. (Supplementary Fig. 7). Collectively, we conclude that VdAve1L2 is an antibacterial effector with a diverged activity spectrum when compared with VdAve1.

**Figure 4.**
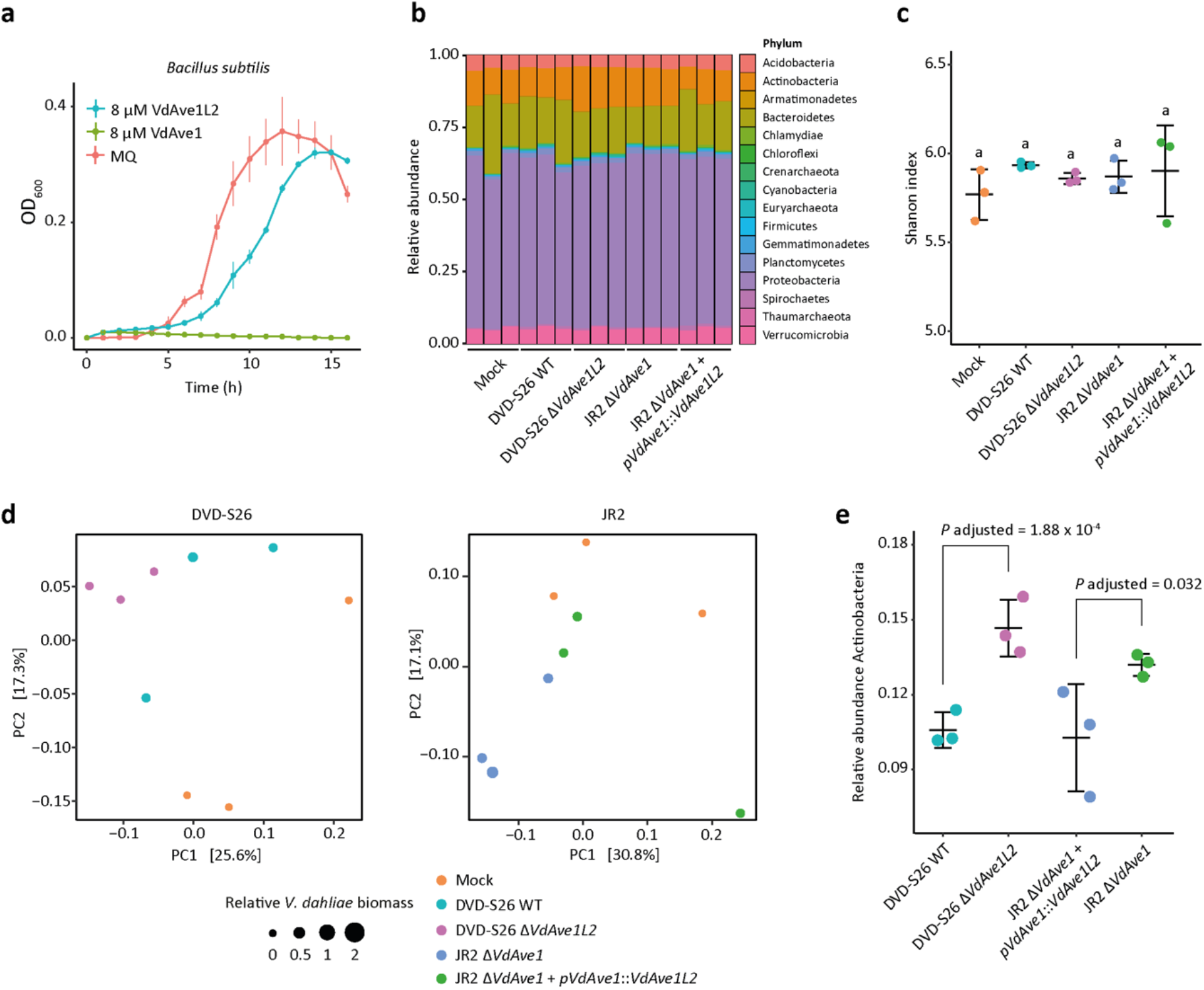
VdAve1L2 impacts Actinobacteria in the tomato root microbiota. **(a)** VdAve1L2 is an antibacterial effector protein. *In vitro* growth of *Bacillus subtilis* is inhibited by VdAve1L2. The previously characterized antibacterial effector VdAve1 (Snelders *et al*., 2020) was included as positive control for bacterial growth inhibition and displays a differential activity compared to VdAve1L2. Ultrapure water (MQ) was included as negative control. Graph displays the average OD_600_ of three biological replicates ± SD. **(b)** Relative abundance of bacterial phyla in tomato root microbiota ten days after inoculation with wild-type *V. dahliae* strain DVD-S26, the corresponding DVD-S26 *VdAve1L2* deletion mutant, a *V. dahliae* strain JR2 *VdAve1* deletion mutant and the corresponding *VdAve1L2* expression mutant as determined by 16S ribosomal DNA profiling. **(c)** *V. dahliae* colonization does not impact α-diversity of tomato root microbiota. The plot displays the average Shannon index ± SD (one-way ANOVA and Tukey’s post-hoc test; p<0.05; N=3). **(d)** Principal coordinate analysis based on Bray-Curtis dissimilarities uncovers separation of root microbiome compositions based on presence of VdAve1L2. **(e)** Differential abundance analysis of bacterial phyla reveals a repression of Actinobacteria in the tomato root microbiota colonized by *V. dahliae* strains that secrete VdAve1L2 (Wald test, N=3).

To determine if VdAve1L2 secretion by *V. dahliae* impacts host microbiota, we performed bacterial community analysis based on 16S ribosomal DNA profiling on tomato roots colonized by *V. dahliae* strain DVD-S26 and the *VdAve1L2* deletion mutant. Furthermore, the *VdAve1* deletion mutant of *V. dahliae* strain JR2 and the corresponding transformant expressing *VdAve1L2* were included. In correspondence with previous observations (Snelders *et al*., 2020), colonization by *V. dahliae* did not dramatically impact the overall composition of bacterial phyla in tomato root microbiota, and also not their α-diversities (Fig. 4b,c). Importantly, however, a principal coordinate analysis based on Bray-Curtis dissimilarities (β-diversity) revealed separation of the bacterial communities based on *V. dahliae* genotype (Fig. 4d), suggesting that secretion of VdAve1L2 impacts root microbiota compositions. Based on pairwise comparisons between the abundances of the bacterial phyla detected in the microbiota in the presence and the absence of VdAve1L2, we identified Actinobacteria as the sole phylum that was significantly suppressed in the microbiota colonized by *V. dahliae* strains secreting VdAve1L2 (Fig. 4e).

To test whether the suppression of Actinobacteria is the direct consequence of antimicrobial activity of VdAve1L2, we incubated representatives of three of the most abundant Actinobacterial families, *Nocardioides, Microbacteriaceae* and *Cryptosporangiaceae*, with VdAve1L2 and monitored their growth *in vitro*. Intriguingly, all tested Actinobacteria displayed higher sensitivity to the effector than most of the other bacteria tested thus far (Fig. 5a), suggesting that Actinobacteria are genuine and direct targets of VdAve1L2 *in planta*.

**Figure 5.**
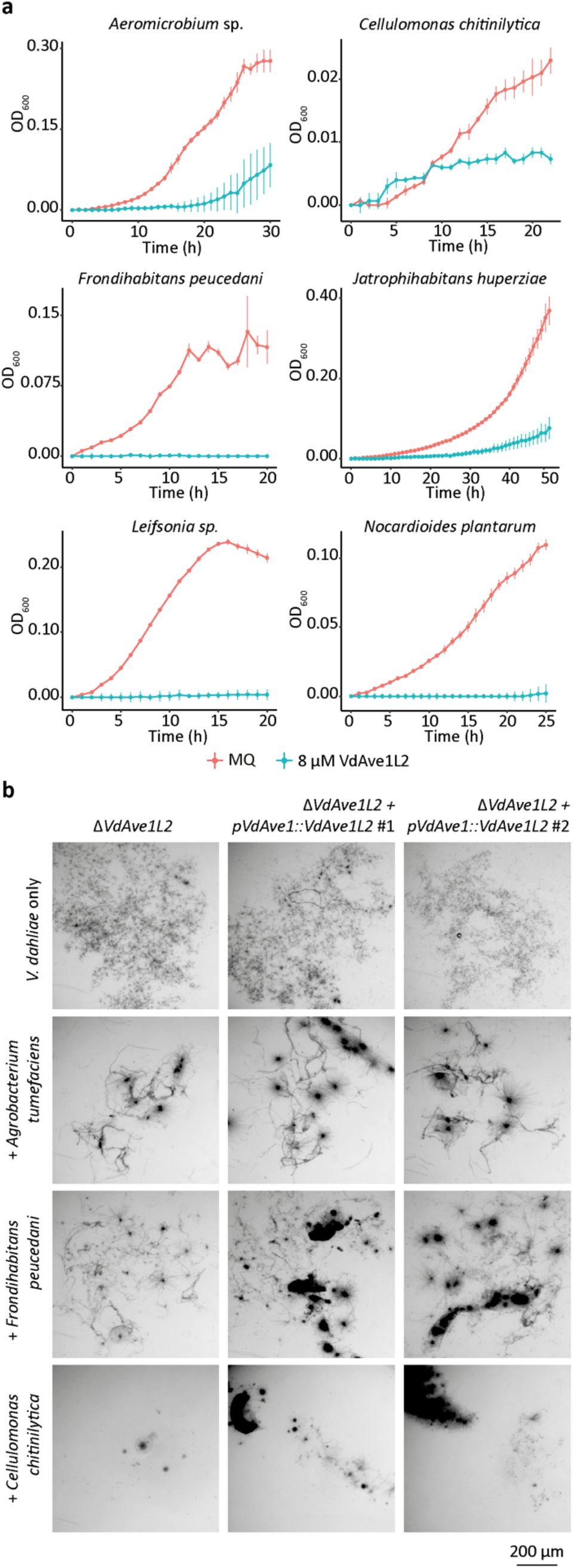
*Verticillium dahliae VdAve1L2* affects antagonistic Actinobacteria. **(a)** Actinobacteria are inhibited by VdAve1L2. Graphs display the average OD_600_ of three biological replicates ±SD. **(b)** VdAve1L2 supports *V. dahliae* growth in the presence of antagonistic Actinobacteria. Representative microscopic pictures displaying the *VdAve1L2* deletion mutant and two mutants expressing *VdAve1L2* under control of the *VdAve1* promoter cultivated for six days in the presence of the *VdAve1L2-*insensitive Proteobacterium *Agrobacterium tumefaciens* and the *VdAve1L2-*sensitive Actinobacteria *Frondihabtians peucedani* and *Cellulomonas chitinilytica*.

Actinobacteria are important players of plant-associated microbial communities, and have repeatedly been assigned roles in disease suppression (Berendsen *et al*., 2018; Chen *et al*., 2020; Lee *et al*., 2021). In accordance with their capacity to produce antimicrobial secondary metabolites, suppression of microbial pathogens by Actinobacteria often involves direct antibiosis, although they have also been implicated in the induction of systemic immunity (Conn *et al*., 2008; Berendsen *et al*., 2018; van Bergeijk *et al*., 2020; Lee *et al*.,2021). To test whether *V. dahliae* exploits VdAve1L2 to compete with Actinobacteria, we co-cultivated the *VdAve1L2* deletion mutant of *V. dahliae* strain DVD-S26 with the VdAve1L2-sensitive Actinobacteria *Frondihabtians peucedani* and *Cellulomonas chitinilytica* and the VdAve1L2-insensitive Proteobacterium *Agrobacterium tumefaciens*. Furthermore, we included transformants of the *VdAve1L2* deletion mutant that express *VdAve1L2* under control of the *VdAve1* promotor that is highly active during *in vitro* growth, in contrast to the VdAve1L2 promoter (Supplementary Fig. 5; Supplementary Fig. 8). As anticipated, secretion of VdAve1L2 failed to counter the antagonistic activity of *A. tumefaciens* and did not promote *V. dahliae* growth when confronted with this bacterium (Fig. 5b). However, *V. dahliae* clearly benefited from VdAve1L2 secretion in competition with both Actinobacteria, as it mediated enhanced fungal growth and development of larger colonies (Fig. 5b). Collectively, our findings suggest that *V. dahliae* secretes VdAve1L2 to antagonize Actinobacteria in the host microbiota.

Next, we aimed to determine the importance of the suppression of Actinobacteria by VdAve1L2 for tomato colonization by *V. dahliae*. Following a previously described protocol (Lee *et al*., 2021), we extracted the root microbiota from tomato plants followed by incubation with the Gram positive bacteria-specific antibiotic vancomycin to affect Actinobacteria. Subsequently, the vancomycin-treated communities and water-treated control communities were allowed to establish on tomato seedlings grown under sterile conditions. Finally, the plants were inoculated with *V. dahliae* strain DVD-S26 and the *VdAve1L2* deletion mutant to determine if manipulation of the Actinobacteria affected *V. dahliae* host colonization and assess the virulence contribution of VdAve1L2. As determined using 16S ribosomal DNA profiling, the tomato plants exposed to the vancomycin-treated microbiota did not harbor a dramatically altered community of bacterial phyla when compared with plants exposed to the water-treated microbiota (Supplementary Fig. 9a). Moreover, the vancomycin treatment did not affect the α-diversity or total abundance of bacteria in the plant microbiota (Supplementary Fig. 9b,c). However, as anticipated, we detected a severe impact of vancomycin treatment on the Actinobacteria community structure (Supplementary Fig. 9d). Moreover, phenotypic assessment revealed markedly increased stunting of tomato plants harboring the vancomycin-treated microbiota when compared with plants containing the water-treated community when inoculated with the *VdAve1L2* deletion mutant, showing disease suppression by Actinobacteria in the water-treated community (Fig. 6a). Quantification of *V. dahliae* biomass in the root microbiomes using real-time PCR confirmed significantly increased colonization by the *VdAve1L2* deletion mutant in the presence of the vancomycin-treated microbiota (Fig. 6b). Importantly, while on plants that were treated with the water-treated community *VdAve1L2* markedly contributes to virulence (Fig. 6a), this virulence contribution is not observed on plants that were treated with the vancomycin-treated community, in line with the hypothesis that the Actinobacteria that are targeted by VdAve1L2 are no longer present in the host microbiota (Fig. 6a). Accordingly, in contrast to the microbiomes with the water-treated microbial community, principal coordinate analysis based on Bray-Curtis dissimilarities (β-diversity) failed to reveal a clear separation of the root microbiomes with the vancomycin-treated community based on their colonization by the different *V. dahliae* strains or mock treatment (Fig. 6c). Moreover, we only detected a VdAve1L2-mediated repression of Actinobacteria genera in plants that received the water-treated communities (Fig. 6d). Likely, treatment with vancomycin limited the abundance of antagonistic Actinobacteria such that interference by VdAve1L2 is no longer required for optimal *V. dahliae* colonization. In conclusion, our findings suggest that Actinobacteria in the tomato root microbiota antagonize host colonization by *V. dahliae*, and that the fungus exploits VdAve1L2 in turn to suppress these antagonists and promote disease.

**Figure 6.**
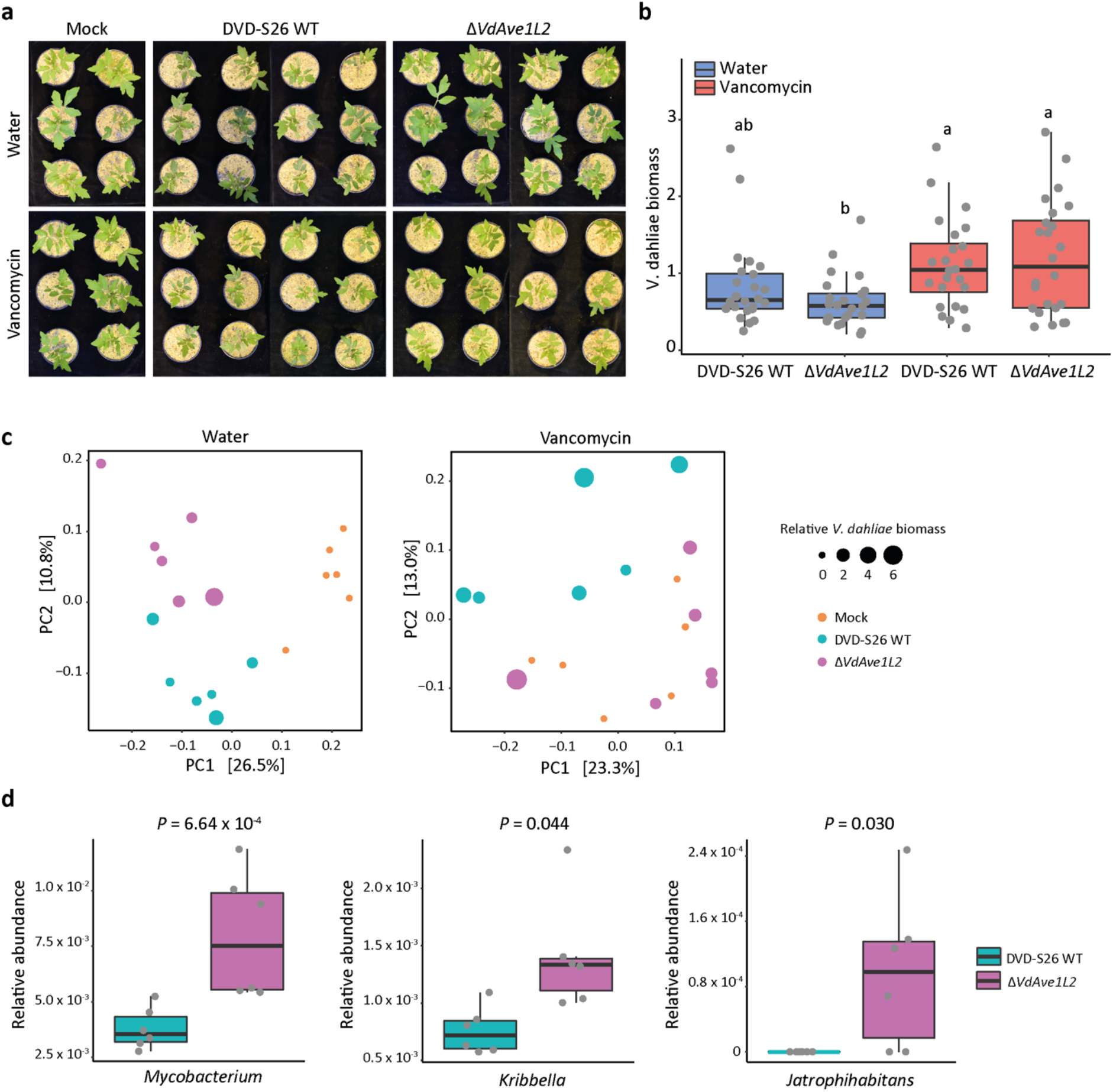
Treatment of tomato root microbiota with vancomycin diminishes the virulence contribution of VdAve1L2. **(a)** Phenotypes of tomato plants harboring vancomycin-treated or water-treated microbial communities infected by wild-type *V. dahliae* and the *VdAve1L2* deletion mutant at 19 days post inoculation. **(b)** Relative *V. dahliae* biomass in tomato stem tissue determined with real-time PCR (one-way ANOVA and Tukey’s post-hoc test; p<0.05; N=24) **(c)** Principal coordinate analysis based on Bray-Curtis dissimilarities uncovers separation of root microbiota compositions in tomato plants harboring water-treated microbial communities, but not vancomycin-treated microbial communities colonized by both *V. dahliae* strains and upon mock treatment. **(d)** Relative abundance of the three Actinobacterial genera that are depleted from the tomato plants harboring the water-treated microbial community colonized by *V. dahliae* WT when compared with colonization by the *VdAve1L2* deletion mutant (Wald test; N=6).

## DISCUSSION

To escape recognition by host immune receptors, microbial plant pathogens rely on the inactivation, loss, or mutation of effectors that become recognized. Unlike many microbial plant pathogens for which allelic effector gene variants escaping recognition have been reported (Armstrong *et al*., 2005; Wu *et al*., 2014), *V. dahliae* was only known to evade effector recognition through purging complete virulence genes, particularly of *VdAve1* and *VdAv2* (de Jonge *et al*., 2012; Chavarro-Carrero *et al*., 2021). Here, we report the discovery of *VdAve1L*,an effector gene with considerable sequence similarity to *VdAve1*, yet that displays extraordinary allelic variation in the *V. dahliae* population, which most likely results from selection pressure imposed by a plant immune receptor.

Functional characterization of the full-length effector variant VdAve1L2 uncovered that this effector, like its homolog VdAve1, exerts antibacterial activity to directly suppress microbial competitors *in planta*. Remarkably, however, VdAve1L2 is exclusively expressed during plant colonization and, in contrast to VdAve1, not expressed *in vitro* or in soil. Moreover, the two effectors display distinct antibacterial activities *in vitro*. Accordingly, we observed that secretion of VdAve1L2 by the race 2 strain DVD-S26, unlike VdAve1 secreted by the race 1 strain JR2, impacts the abundance of Actinobacteria and not of Sphingomonadales in tomato. VdAve1L2 and VdAve1 thus functionally diverged from each other. So far, the mode of action of VdAve1 remains unclear and, accordingly, it is unclear how VdAve1L2 functionally diverged from VdAve1 to target Actinobacteria rather than Sphingomonadales.

Actinobacteria represent a core phylum that is found in virtually any plant grown in any environment. Several Actinobacterial species were shown to fulfill beneficial roles in plant holobionts, such as suppression of plant diseases (Berendsen *et al*., 2018; Chen *et al*., 2020; Lee *et al*., 2021). Additionally, Actinobacteria are keystone taxa that impact and benefit microbial community structures in plants (Carlström *et al*., 2019; Gómez-Pérez *et al*., 2022).

Considering these beneficial traits, it is not surprising that members of this phylum are targeted by microbial plant pathogens to weaken plant holobionts. Interestingly, the oomycete Arabidopsis pathogen *Albugo candida* was recently reported to deposit several antibacterial effector proteins in the leaf apoplast (Gómez-Pérez *et al*., 2022). Interestingly, some of these effectors impact growth of Actinobacterial keystone taxa of the Arabidopsis phyllosphere *in vitro*, suggesting that the suppression of Actinobacteria in host microbiota might be a strategy adopted by diverse microbial plant pathogens. These findings furthermore support the hypothesis that effector-mediated manipulation of host microbiota communities may be a widely deployed strategy of plant pathogens to support host colonization (Snelders *et al*., 2022).

VdAve1 is recognized by the tomato immune receptor SlVe1, encoded by a gene in a locus that also encodes the highly similar orphan receptor SlVe2 (Fradin *et al*., 2014). Considering the similarity between *VdAve1* and *VdAve1L*, we tested if SlVe2 was able to recognize any of the current *VdAve1L* alleles or their putative progenitors, but none of those evoked a detectable hypersensitive response upon overexpression in combination with SlVe2. However, if recognition took place in tomato, it may equally well have been mediated by any other putative immune receptor encoded in the tomato genome. Perhaps even more likely, recognition may also have occurred in any of the hundreds of other *V. dahliae* hosts. Importantly, Actinobacteria are ubiquitously present in a wide diversity of plants, and thus *V. dahliae* is likely to benefit from the antibacterial activity of VdAve1L2 in plant species beyond tomato.

We previously showed that *VdAve1* was horizontally acquired from plants, where the abundantly present homologs are generally annotated as plant natriuretic peptides (PNPs) (de Jonge *et al*., 2012). The fact that some of the sequenced *V. dahliae* isolates carry both *VdAve1* and *VdAve1L* raises the question if both genes have been introduced by two separate horizontal gene transfer (HGT) events, or whether only a single HGT event took place that was followed by gene duplication and divergence. Although we have tried to resolve what scenario is most likely, our (phylogenetic) analyses rendered inconclusive results. Hence, at present the exact relationship between VdAve1 and VdAve1L remains unclear. Nevertheless, with VdAve1L2 we here reported the characterization of the fourth *V. dahliae* effector protein that acts in microbiota manipulation (Snelders *et al*., 2020, 2021), which following VdAve1 (de Jonge *et al*., 2012), is most likely the second microbiota-manipulating effector secreted by *V. dahliae* that is recognized by a plant immune receptor. In light of the view that the microbiota constitutes an extrinsic layer of the plant immune system (Dini-Andreote, 2020), microbiota-manipulating effectors target a critical immune component of the host, and thus constitute a relevant target for surveillance by plants to mediate timely pathogen detection, in a similar fashion as effector proteins that interfere with intrinsic immune components are perceived. As a consequence of recognition, microbial plant pathogens need to mutate, purge or inactivate their microbiota-manipulating effector proteins to escape host recognition, which leads to pathogen races with divergent suites of antimicrobial effectors. A possibility for the more effective use of microbial biocontrol agents could be to base their selection on the genotype of a plant pathogen, for instance by selecting antagonists that are insensitive to the activity of a specific (lineage-specific) effector. Conversely, in case a resistance gene has been described to recognize a microbiota-manipulating effector protein, the application of a strong antagonistic biocontrol agent that is sensitive towards the activity of the corresponding effector can be considered. In this manner, a strong selection pressure is exerted to retain that particular effector gene in the pathogen, which may contribute to enhanced durability of the resistance in turn. In this manner, the further identification and characterization of microbiota-manipulating effectors secreted by microbial plant pathogens may aid in the development of more sophisticated, and perhaps more successful, biocontrol strategies.

## ACKNOWLEDGEMENTS

Y.S., G.L.F. and D.E.T. acknowledge PhD fellowships from the China Scholarship Council (CSC), Coordination for the Improvement of Higher Education Personnel (CAPES) from the federal government of Brazil, and the Consejo Nacional de Ciencia y Tecnología de México, respectively. The authors thank Bert Essenstam and Pauline Sanderson (Unifarm) for excellent plant care, as well as José Espejo Valle-Inclan for assistance with the Oxford Nanopore MinION sequencing. B.P.H.J.T acknowledges funding by the Alexander von Humboldt Foundation in the framework of an Alexander von Humboldt Professorship endowed by the German Federal Ministry of Education and Research is furthermore supported by the Deutsche Forschungsgemeinschaft (DFG, German Research Foundation) under Germany’s Excellence Strategy – EXC 2048/1 – Project ID: 390686111 and by the Research Council for Earth and Life Science (ALW) of the Netherlands Organization for Scientific Research (NWO). The authors declare no conflict of interest exists.

## AUTHOR CONTRIBUTIONS

N.C.S., J.C.B., and B.P.H.J.T. conceived the project. N.C.S., J.C.B., Y.S., N.S., G.L.F., H.R., G.C.M.B, D.E.T., L.F., M.F.S. and B.P.H.J.T designed and performed the experiments. N.C.S., J.C.B., Y.S., N.S., G.L.F., H.R., G.C.M.B, D.E.T., L.F., M.F.S. and B.P.H.J.T. analyzed the data. N.C.S., J.C.B., G.L.F. and B.P.H.J.T. wrote the manuscript. All authors read and approved the final manuscript.

## DATA AVAILABLILITY

The 16S profiling data have been deposited in the NCBI GenBank database under BioProject PRJNA742137.

## Notes

### Competing Interest Statement

The authors have declared no competing interest.

