## Supplementary Data for "A highly polymorphic effector protein promotes fungal virulence through suppression of plant-associated Actinobacteria"

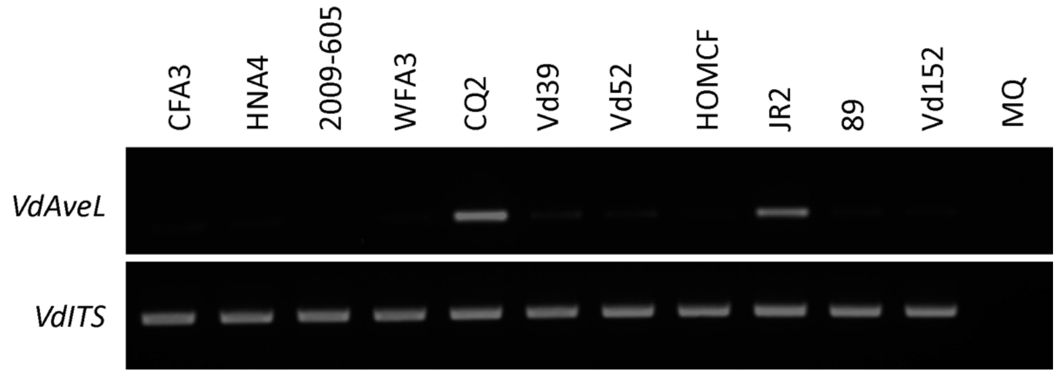


**Supplementary Figure 1. Presence/absence of *VdAve1L* in *V. dahliae* strains as determined by PCR.** PCR confirms the absence of *VdAve1L* in CFA3, HNA4, 2009-605, WFA3, Vd39, HOMCF, 89 and Vd152. The *VdAve1L*-containing strains CQ2 and JR2 were included as positive controls. *VdITS* was included as genomic DNA control.

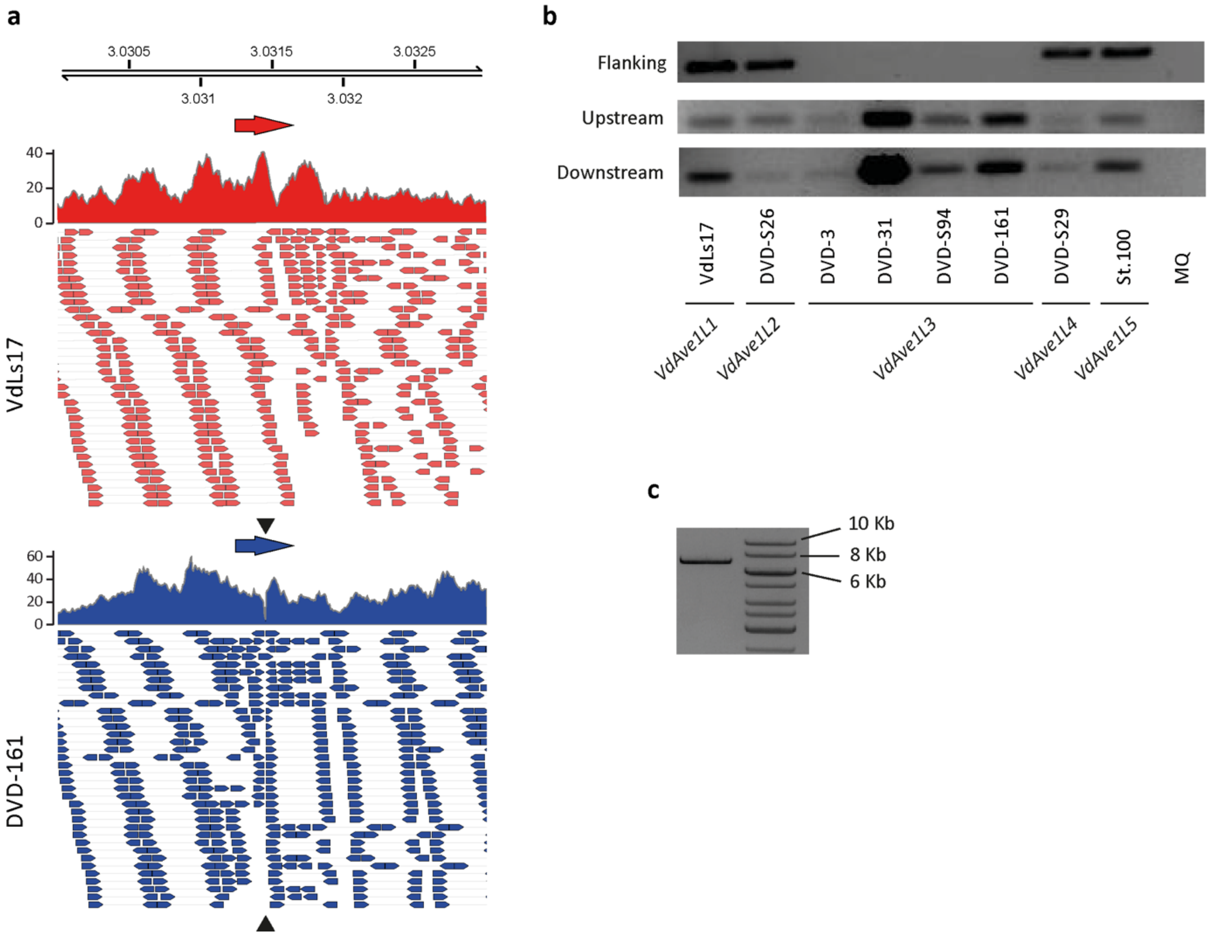


**Supplementary Figure 2. *VdAve1L3* is disrupted by a transposable element. (a)** Read coverage of *V. dahliae* strain VdLs17 (*VdAve1L1*) and DVD-161 (*VdAve1L3*) across the genomic region surrounding *VdAve1L* of the reference genome of JR2. The red and blue arrows indicate the location of *VdAve1L* and the black arrows indicate the discontinuity of *VdAve1L3*. **(b-c)** PCR confirms discontinuity in *VdAve1L3*. **(b)** Amplicons obtained by PCR using primers that allow amplification of the regions flanking or spanning the site where the discontinuity in *VdAve1L3* occurs. Regular elongation time did not yield an amplicon for the region spanning the discontinuity site in strains with the *VdAve1L3* allele. **(c)** Extended elongation time allows amplification of the region spanning the discontinuity site in *VdAve1L3* of *V. dahliae* strain DVD-3. The obtained amplicon size indicates that *VdAve1L3* contains an insert of approximately 7 Kb. Sequence analysis of the amplicon revealed that *VdAve1L3* is interrupted by a long terminal repeat retrotransposon that is classified as VdLTRE3 (Faino *et al.*, 2016).

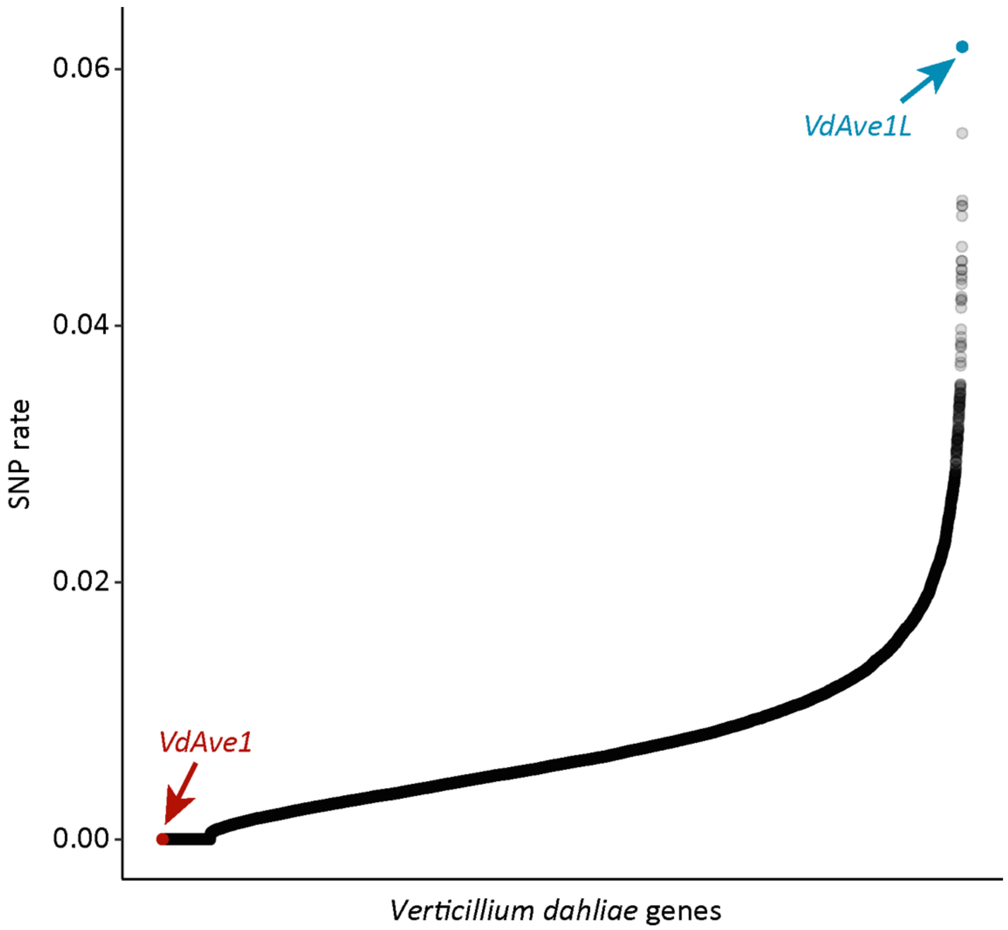


**Supplementary Figure 3. The *VdAve1L* allele displays extraordinary sequence variation.** Each dot displays the detected SNP rate in the coding sequence of a single gene in the genome of *V. dahliae* strain JR2, with *VdAve1L* displaying the highest variation. The previously identified *V. dahliae* (a)virulence gene *VdAve1*, that like *VdAve1L* also displays presence-absence variation in the *V. dahliae* strains, shows no sequence variation.

```
VdAve1L4    ATGAAGCTTTCTACGCTTGGAGCCCTCATTTCATTGACTCCACTAGTCACTGCCGAAGTA    60
VdAve1L5    ATGAAGCTTTCTACGCTTGGAGCCCTCATTTCATTGACTCCACTAGTCACTGCCGAAGTA    60
VdAve1L6    ATGAAGCTTTCTACGCTTGGAGCCCTCATTTCATTGACTCCACTAGTCACTGCCGAAGTA    60
VdAve1L3    ATGAAGCTTTCTACGCTTGGAGCCCTCATTTCATTGACTCCACTAGTCACTGCCGACGTA    60
VdAve1L1    ATGAAGCTTTCTACGCTTGGAGCCCTCATTTCATTGACTCCACTAGTCACTGCCGACGTA    60
VdAve1L2    ATGAAGCTTTCTACGCTTGGAGCCCTCATTTCATTGACTCCACTAGTCACTGCCGACGTA    60
            ******************************************************** ***

VdAve1L4    GGGACCGCATCCTAGTATCCTCCACCCTACACTCCTAATTATTGCCACATCGATGCGAGC    120
VdAve1L5    GGGACCGCATCCTGGTATCCTCCACCCTACACTCCTAATTATTGCCACATCGATGCGAGC    120
VdAve1L6    GGGACCGCATCCTGGTATCCTCCACCCTACACTCCTAATTATTGCCACATCGATGCGAGC    120
VdAve1L3    GGGACCGCATCCTAGTATCCTCCACCCTACACTCCTAATTATTGCTATATCGATGCGAGC    120
VdAve1L1    GGGACCGCATCCTAGTATCCTCCACCCTACACTCCTAATTATTGCTACATCGATGCGAGC    120
VdAve1L2    GGGACCGCATCCTGGTATCCTCCACCCTACACTCCTAATTATTGCTACATCGATGCGAGC    120
            ************* ******************************* * ************

VdAve1L4    AGATTCCCCAATGATGTTCATATTGTTTCTTTTTCAGAAGAACTCATGCAAGGCGGTGCT    180
VdAve1L5    AGATTCCCCAATGATGTTCATATTGTTTCTTTTTCAGAAGAACTCATGCAAGGCGGTGCT    180
VdAve1L6    AGATTCCCCAATGATGTTCATATTGTTTCTTTTTCAGAAGAACTCATGCAAGGCGGTGCT    180
VdAve1L3    AGATTCCCCGATAATAGTTATATTGTTTCTTTTTCAGAAGAACTCATGTAAGGCGGTGCT    180
VdAve1L1    AGATTCCCCGATAATAGTTATATTGTTTCTTTTTCAGAAGAACTCATGCAAGGCGGTGCT    180
VdAve1L2    AGATTCCCCGATGATGGTTATATTGTTTCTTTTTCAGAAGAACTCATGCAAGGCGGTGCT    180
            ********* ** **  * ***************************** ***********

VdAve1L4    GTCTATGGTATGAGGTACTTAGTTAAATGCATTAGTCTTAAAGAGAGTAAATCGGGCTGT    240
VdAve1L5    GTCTATGGTATGAGGTACTTAGTTAAATGCATTAGTCTTAATGAGAGTAAATCGGGCTGT    240
VdAve1L6    ATCTATGGTATGAGGTACTTAGTTAAATGCATTAGTCTTAATGAGAGTAAATCGGGCTGT    240
VdAve1L3    GTCTATAGTGAGAGTTACTTAGTTACATATATGAGTCTTAATGAGAGGAAATCGGGCTAT    240
VdAve1L1    GTCTATGGTGAGAGTTACTTAGTTAGATGCATTAGTCTTAATGAGAGGAAATCGGGCTGT    240
VdAve1L2    GTCTGTGGTGAGAGTTACTTAGTTACATGCATGAGTCTTAATGAGAGGAAATCGGGCTGT    240
             *** * **  *** ********** **  ** ******** ***** ********** *

VdAve1L4    AGGCATACTGAACCGATTGAAGTGATGGTAGTTGATCAAGGCAAGGAAGGTACTACTCAT    300
VdAve1L5    AGGCATACTGAACCGATTGAAGTGATGGTAGTTGATCAAGGCAAGGAAGGTACTACTCAT    300
VdAve1L6    AGGCATACTGAACCGATTGAAGTGATGGTAGTTGATCAAGGCAAGGAAGGTACTACTCAT    300
VdAve1L3    AGGCATACTGAACCGATTGAAGTGATAGTAGTTGATCAAGGCAAGGAAGGTACTACTCAT    300
VdAve1L1    AGGCATACTGAACCGATTGAAGTGATGGTAGTTGATCAAGGCAAGGAAGGTACTACTCAT    300
VdAve1L2    AGGCATACTGAACCGATTGAAGTGATGGTAGTTGATCAAGGCAAGGAAGGTACTACTCAT    300
            ************************** *********************************

VdAve1L4    CAAGGACATAAACCGAAAATGGTTCTTTCTAAGGAGGCTTATATTGCTCTTATTAAACAG    360
VdAve1L5    CAAGGACATAAACCGAAAATGGTTCTTTCTAAGGAGGCTTATATTGCTCTTGTTAAACAG    360
VdAve1L6    CAAGGACATAAACCGAAAATGGTTCTTTCTAAGGAGGCTTATATTGCTCTTGTTAAATAG    360
VdAve1L3    CAAGGACATAAACCGAAAATGGTTCTTTCTAAGGAGGCTTATATTGCTCTTGTTAAATAG    360
VdAve1L1    CAAGGACATGAACCGAAAATGGTTCTTTCTAAGGAGGCTTATATTGCTCTTGTTAAACAG    360
VdAve1L2    CAAGGACATAAACCGAAAATGGTTCTTTCTAAGGAGGCTTATATTGCTCTTGTTAAACAG    360
            ********* ***************************************** ***** **

VdAve1L4    TATATTAAGCATTCGTTCATTCAAGTTGAGTATAGAATAAAATAA    405
VdAve1L5    TATGTTAAGCATTCGTTCATTCAAGTTGAGTATAGAATAAAATAA    405
VdAve1L6    TATGTTAAGCATTCGTTCATTCAAGTTGAGTATAGAATAAAATAA    405
VdAve1L3    TATGTTAAGCATTCGTTCATTTAAGTTGAGTATAGAAAAAAATAA    405
VdAve1L1    TATATTAAGCATTCGTTCATTCAAGTTGAGTATAGAAAAAAATAA    405
VdAve1L2    TATGTTAAGCATTCGTTCATTCAAGTTGAGTATAGAAAAAAATAA    405
            *** ***************** *************** *******
```

**Supplementary Figure 4. SNP locations in the CDS of the *VdAve1L* alleles.** Sequence alignment of the coding sequences of the VdAve1L alleles when premature stop codons and/or transposon insertions are ignored. The SNPs, all non-synonymous, are highlighted in red.

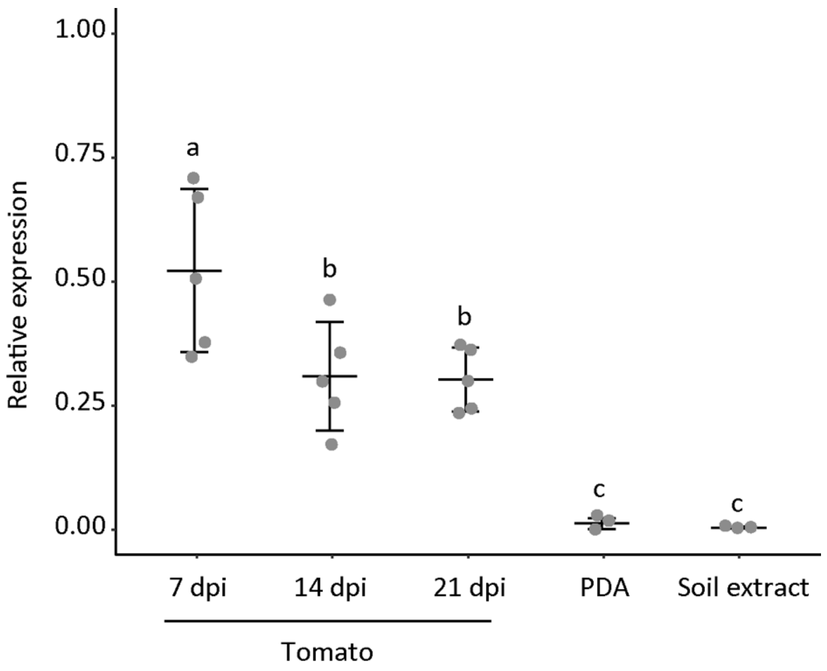


**Supplementary Figure 5. *VdAve1L2* is expressed by *Verticillium dahliae* during tomato colonization.** The graph displays expression of *VdAve1L2* relative to *VdGAPDH* during colonization of tomato at 7, 14 and 21 days post inoculation (N=5), growth on potato dextrose agar at 5 days of cultivation (N=3), and growth in soil extract at 5 days of cultivation (N=3). *V. dahliae* expresses *VdAve1L2* during tomato colonization, but not during growth on PDA or in soil extract. Different letter labels indicate significant differences (one-way ANOVA and Tukey's post-hoc test; $p<0.05$).

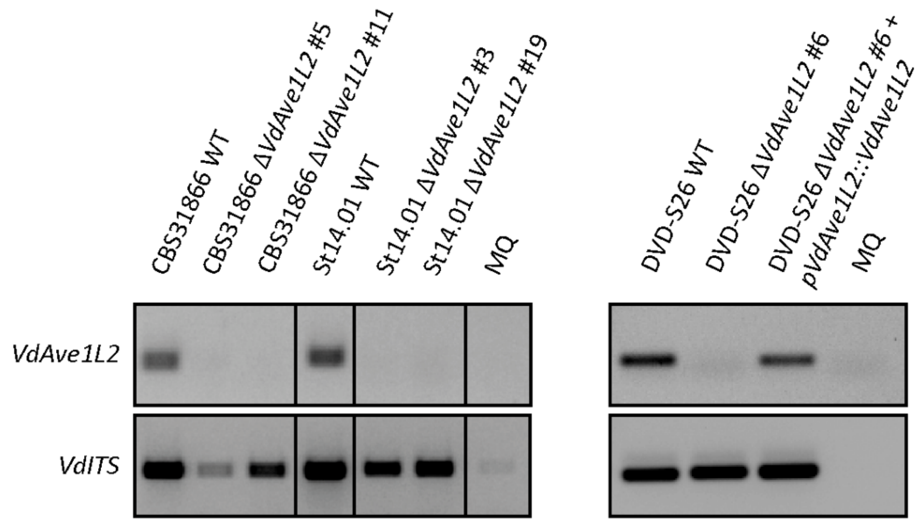


**Supplementary Figure 6. Verification of *VdAve1L2* deletion and complementation mutants.** Presence/absence of *VdAve1L2* in deletion and complementation mutants was verified by PCR, *VdITS* was included as genomic DNA control.

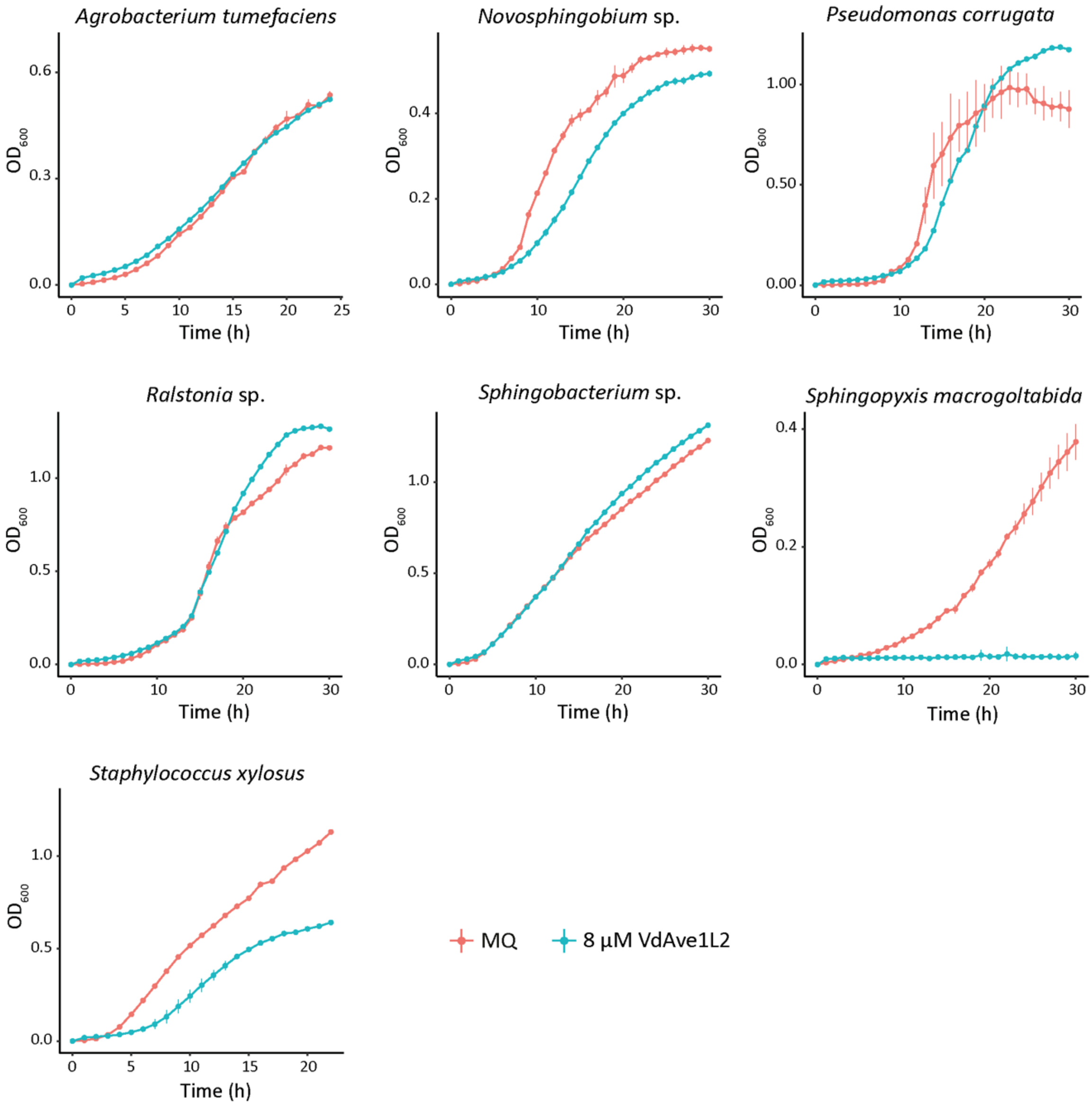


**Supplementary Figure 7. VdAve1L2 is an antibacterial effector protein.** Growth of plant-associated bacterial isolates *in vitro* in presence of VdAve1L2 or ultrapure water (MQ) as negative control. Graphs display the average $OD_{600}$ of three biological replicates ± SD.

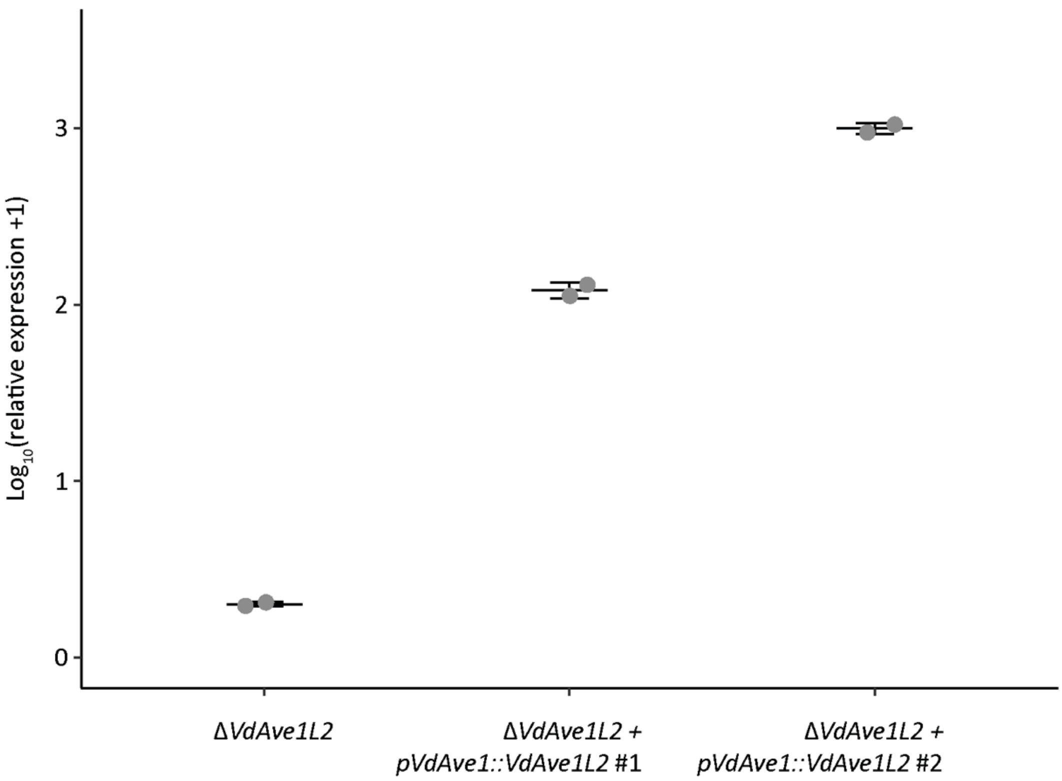


**Supplementary Figure 8. Expression of *VdAve1L2* in *V. dahliae* mutants.** Expression of *VdAve1L2* in two *V. dahliae* DVD-S26 mutants expressing *VdAve1L2* under control of the *VdAve1* promoter relative to the DVD-S26 *VdAve1L2* deletion mutant after five days of cultivation in 0.05x PDB (N=2).

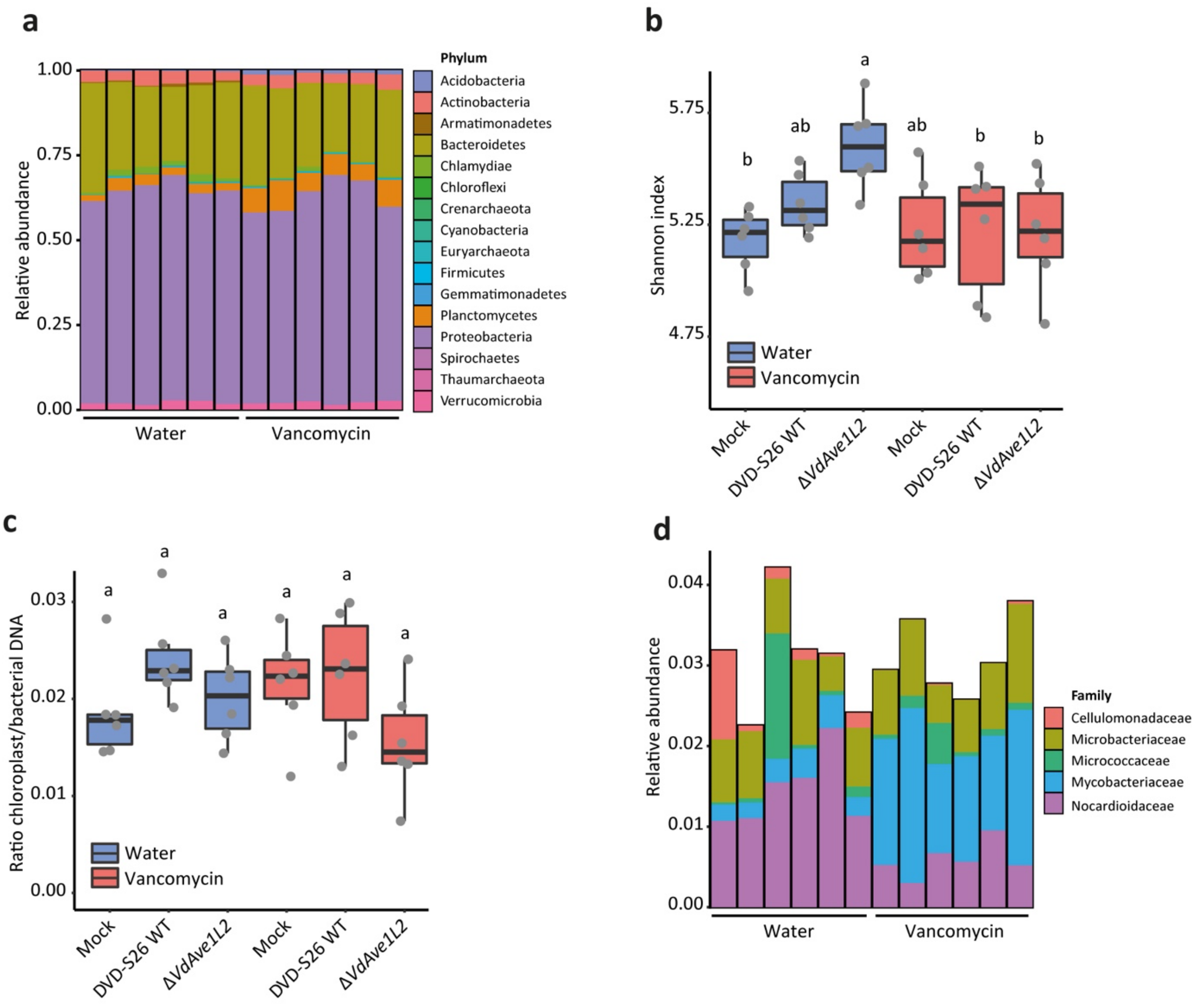


**Supplementary Figure 9. Metagenomic characterization of tomato root microbiota with water-treated and vancomycin-treated microbial communities. (a)** Relative abundance of bacterial phyla in the root microbiomes of mock-inoculated tomato plants that received the water-treated or vancomycin-treated microbial communities nineteen days post transplantation to river sand with Hoagland solution. **(b)** Vancomycin treatment and *V. dahliae* colonization does not dramatically impact the α-diversity of tomato root microbiomes. **(c)** Vancomycin treatment did not impact the total abundance of bacteria in the tomato root microbiota. Boxplots display the ratio between the plant chloroplast reads and bacterial reads obtained in the 16S rDNA profiling. **(d)** Vancomycin treatment did not impact the relative abundance of the Actinobacteria phylum, but severely impacted Actinobacterial community structures. Bar plot displays the relative abundance of Actinobacteria families in the root microbiomes of the mock-inoculated tomato plants that received the water-treated or vancomycin-treated microbial communities.

**Supplementary Table 1.** List of primers used in this study.

| Name | Oligonucleotide Sequence (5'-3') | Description |
|---|---|---|
| Ave1L-Break-F | TCGATGCGAGCAGATTCC | Primers spanning discontiguity |
| Ave1L-Break-R | GAGTAGTACCTTCCTTGCCTTGA | Primers spanning discontiguity |
| Ave1L-Up-F | CTACGCTTGGAGCCCTCAT | Primers upstream of discontiguity |
| Ave1L-Up-R | AGGAGTGTAGGGTGGAGGAT | Primers upstream of discontiguity |
| Ave1L-Down-F | GGCATACTGAACCGATTGAAG | Primers downstream of discontiguity |
| Ave1L-Down-R | CTCCTTAGAAAGAACCATTTTCG | Primers downstream of discontiguity |
| VdLs17del.stop_F | CGCATCCTGGTATCCTCCACC | Replace stop codon in *VdAve1L1* and *VdAve1L4* |
| VdLs17del.stop_R | TGGAGGATACCAGGATGCGGT | Replace stop codon in *VdAve1L1* and *VdAve1L4* |
| Topo_Ave1L_F | CACCATGAAGCTTTCTACGCTTGGAG | Combined with primer VdLs17del.stop_Rv |
| Topo_St14.01_R | TTATTTTTTTCTATACTCAACTTGAATGAAC | Combined with primer VdLs17del.stop_Fw |
| Topo_St.100_R | TTATTTTATTCTATACTCAACTTGAATGAAC | Combined with primer VdLs17del.stop_Fw |
| Ave1L2-LB-F3 | ggtcttaauTTTTATCTCTCCCTTCCTCTTATCT | 1500 bp fragment upstream of *VdAve1L2* |
| Ave1L2-LB-R3 | ggcattaauTTTTTAAGCCTTTCTAGCTTATTCTT | 1500 bp fragment upstream of *VdAve1L2* |
| Ave1L2-RB-F3 | ggacttaauGCTATCTTCACGAGAGCAGAGT | 1500 bp fragment downstream of *VdAve1L2* |
| Ave1L2-RB-R3 | gggtttaauTTGCGCGTTTATATATTCTTATCTT | 1500 bp fragment downstream of *VdAve1L2* |
| Ave1L2_1kb_Up_attB2r_F | ggggacagctttcttgtacaaagtggaaGCTATAAGTCCTTTAGTGCCTATCC | Amplification of *VdAve1L2* with 1000 bp upstream and downstream |
| Ave1L2_1kb_down_attB3_R2 | ggggacaactttgtataataaagttgtTATAGGTCTAGAGGAAGTCTAAGCG | Amplification of *VdAve1L2* with 1000 bp upstream and downstream |
| Ave1L2_gen_F | CTCCACCCTACACTCCTAATTATT | Genotyping of *VdAve1L2* in *V. dahliae* strains |
| Ave1L2_gen_R | AGTATGCCTACAGCCCGATT | Genotyping of *VdAve1L2* in *V. dahliae* strains |
| Ave1L2_qPCR_F | CTTTCTACGCTTGGAGCCCT | Determination of *VdAve1L2* gene expression |
| Ave1L2_qPCR_F | ACCGCCTTGCATGAGTTCTT | Determination of *VdAve1L2* gene expression |
| ITS1-F | AAAGTTTAATGGTTCGCTAAGA | *V. dahliae* biomass quantification |
| StVe1-R | CTTGGTCATTTAGAGGAAGTAA | *V. dahliae* biomass quantification |
| SIRub_F | GAACAGTTTCTCACTGTTGAC | *S. lycopersicum* RuBisCO, real-time PCR |
| SIRub_R | CGTGAGAACCATAAGTCACC | *S. lycopersicum* RuBisCO, real-time PCR |
